# MVA Vector Vaccines Inhibit SARS CoV-2 Replication in Upper and Lower Respiratory Tracts of Transgenic Mice and Prevent Lethal Disease

**DOI:** 10.1101/2020.12.30.424878

**Authors:** Ruikang Liu, Jeffrey L. Americo, Catherine A. Cotter, Patricia L. Earl, Noam Erez, Chen Peng, Bernard Moss

**Affiliations:** Laboratory of Viral Diseases, National Institute of Allergy and Infectious Diseases, National Institutes of Health, Bethesda MD 20892 USA

## Abstract

Replication-restricted modified vaccinia virus Ankara (MVA) is a licensed smallpox vaccine and numerous clinical studies investigating recombinant MVAs (rMVAs) as vectors for prevention of other infectious diseases have been completed or are in progress. Two rMVA COVID-19 vaccine trials are at an initial stage, though no animal protection studies have been reported. Here, we characterize rMVAs expressing the S protein of CoV-2. Modifications of full length S individually or in combination included two proline substitutions, mutations of the furin recognition site and deletion of the endoplasmic retrieval signal. Another rMVA in which the receptor binding domain (RBD) flanked by the signal peptide and transmembrane domains of S was also constructed. Each modified S protein was displayed on the surface of rMVA-infected human cells and was recognized by anti-RBD antibody and by soluble hACE2 receptor. Intramuscular injection of mice with the rMVAs induced S-binding and pseudovirus-neutralizing antibodies. Boosting occurred following a second homologous rMVA but was higher with adjuvanted purified RBD protein. Weight loss and lethality following intranasal infection of transgenic hACE2 mice with CoV-2 was prevented by one or two immunizations with rMVAs or by passive transfer of serum from vaccinated mice. One or two rMVA vaccinations also prevented recovery of infectious CoV-2 from the lungs. A low amount of virus was detected in the nasal turbinates of only one of eight rMVA-vaccinated mice on day 2 and none later. Detection of subgenomic mRNA in turbinates on day 2 only indicated that replication was abortive in immunized animals.

**Significance:** Vaccines are required to control COVID-19 during the pandemic and possibly afterwards. Recombinant nucleic acids, proteins and virus vectors that stimulate immune responses to the CoV-2 S protein have provided protection in experimental animal or human clinical trials, though questions remain regarding their ability to prevent spread and the duration of immunity. The present study focuses on replication-restricted modified vaccinia virus Ankara (MVA), which has been shown to be a safe, immunogenic and stable smallpox vaccine and a promising vaccine vector for other infectious diseases and cancer. In a transgenic mouse model, one or two injections of recombinant MVAs that express modified forms of S inhibited CoV-2 replication in the upper and lower respiratory tracts and prevented severe disease.

## Introduction

Recombinant DNA methods have revolutionized the engineering of vaccines against microbial pathogens, thereby creating opportunities to control the current SARS CoV-2 pandemic (1). The main categories of recombinant vaccines are protein, nucleic acid (DNA and RNA), virus vectors (replicating and non-replicating) and genetically modified live viruses. Each approach has distinctive advantages and drawbacks with regard to manufacture, stability, cold-chain requirements, mode of inoculation, and immune stimulation. Recombinant proteins have been successfully deployed as hepatitis B, papilloma, influenza and varicella Zoster virus vaccines (2–5). DNA vaccines have been licensed to protect horses from West Nile virus and salmon from infectious hematopoietic necrosis virus (6, 7), though none are in regular human use. Recently developed mRNA vaccines received emergency approval for COVID-19 and are in pre-clinical development for other infectious diseases (8). At least 12 virus vector vaccines based on adenovirus, fowlpox virus, vaccinia virus (VACV) and yellow fever virus have veterinary applications, but so far only an attenuated yellow fever vectored Dengue and a chimeric Japanese encephalitis virus vaccine have been marketed for humans (9), though numerous clinical trials particularly with attenuated adenovirus and VACV are listed in ClinicalTrials.gov.

A variety of recombinant approaches utilizing the spike (S) protein as immunogen are being explored to quell the SARS CoV-2 pandemic (10). Vaccines based on mRNA and adenovirus have demonstrated promising results in animal models as well as in clinical trials and some have already received emergency regulatory approval (11–14). Other vaccines, including ones based on vesicular stomatitis virus (15), an alphavirus-derived replicon RNA (16), and an inactivated recombinant New Castle Disease virus (17) have shown protection in animal models. Immunogenicity in mice was found for a modified VACV Ankara (MVA)-based CoV-2 vaccine, (18), but animal protection studies have not yet been reported. However, protection has been obtained with related MVA-based SARS CoV-1 and MERS in animals (19–22) and a MVA MERS vaccine was shown to be safe and immunogenic in a phase 1 clinical trial (23).

Experiments with virus vectors for vaccination were carried out initially with VACV (24, 25), providing a precedent for a multitude of other virus vectors (9). The majority of current VACV vaccine studies employ the MVA strain, which was attenuated by passage more than 500 times in chicken embryo fibroblasts (CEF) during which numerous genes were deleted or mutated resulting in an inability to replicate in human and most other mammalian cells (26). Despite the inability to complete a productive infection, MVA is capable of highly expressing recombinant genes and inducing immune responses (27, 28). MVA is a licensed smallpox vaccine and numerous clinical studies of recombinant MVA (rMVA) vectors are in progress or have been completed and two for COVID-19 are in the recruiting phase (ClinicalTrials.gov). Here, we show that one or two immunizations with rMVAs expressing SARS CoV-2 spike proteins elicit strong neutralizing antibody responses, induce CD8+ T-cells and protect susceptible transgenic mice against a lethal intranasal challenge with CoV-2 virus, supporting clinical testing of related rMVA vaccines.

## Results

### Construction of rMVAs and expression of S proteins

The full-length CoV-2 S protein contains 1273 amino acids (aa) comprising a signal peptide (aa 1-13), the S1 receptor binding subunit (aa 14-685) and the S2 membrane fusion subunit (aa 686-1273). A panel of rMVAs with names in italics including *WT* expressing unmodified CoV-2 S (GenBank: QHU36824) and modified forms of the S protein with C-terminal FLAG tags were engineered (Fig. 1). The rMVAs with modified versions of full-length S include *2P* with two proline substitutions (K_986_P, V_987_P) intended to stabilize the prefusion conformation (11, 29–31), *Δfurin* with perturbation of the furin recognition site (RRAR_682-685_GSAS) to prevent cleavage of S, *ΔERRS* with deletion of the last 19 aa including the endoplasmic reticulum retrieval signal and *Tri* with a combination of all three modifications. *RBD*, another rMVA, contains the receptor binding domain (RBD) and contiguous sequences preceded by the S signal peptide and followed by the transmembrane domain of S. The latter were added because a previous study with an unrelated protein demonstrated that membrane anchoring strongly enhances immunogenicity of a VACV vector (32) and also enhanced immunogenicity of CoV-2 S expressed by an mRNA vaccine (11). The additional rMVAs, *D_614_*G and a *2P* version, express S with amino acid changes of a recently emerged strain (33). VACV transcription termination signals that could reduce early expression (34) and runs of four or more consecutive Gs or Cs that could accelerate the occurrence of deletions (35) were altered by making silent mutations.

**Fig. 1.**
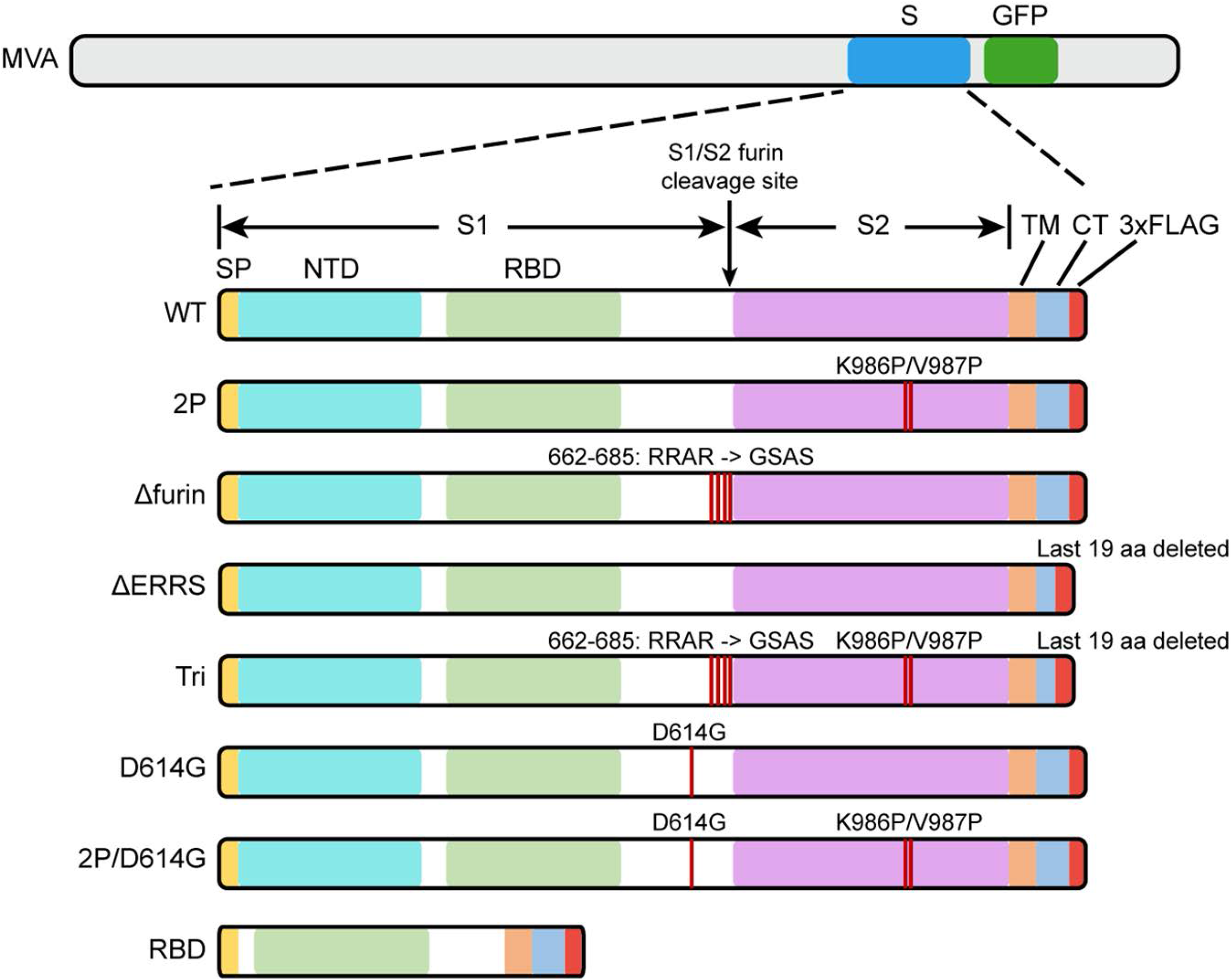
Diagrams of rMVAs. Top shows approximate locations of CoV-2 spike protein (S) and green fluorescent protein (GFP) ORFs within rMVA. Modifications of S ORF are shown below with names of constructs on the left. Abbreviations: SP, signal peptide; NTD, N-Terminal domain; TM, transmembrane domain; CT, C-terminal domain; RBD, receptor binding domain; 3xFLAG, 3 tandem copies of FLAG epitope tag.

DNA encoding the modified S open reading frames (ORFs) were inserted into the pLW44 transfer vector (36), which provides a VACV early/late promoter and flanking sequences that enable directed recombination into a non-essential region of the MVA genome and selection of plaques in CEF by green fluorescence. After multiple rounds of plaque purification, the sequences of WT and modified S ORFs were confirmed and the viruses were expanded in CEF and purified by sedimentation through a sucrose cushion.

Since CoV-2 vaccines are intended for humans, in which replication of MVA is restricted, HeLa cells were used to evaluate expression of the S proteins during a single round of infection. Cell lysates were analyzed by SDS-polyacrylamide gel electrophoresis and Western blotting. Full-length S proteins of ~180 kDa or a ~50 kDa shortened version in the case of *RBD*, were detected by antibodies to the RBD (Fig. 2A) and to the C-terminal FLAG tag (Fig. 2B). The anti-RBD antibody also recognized S1, formed by cleavage of full-length S, from lysates of cells infected with *WT, D_614_*G and their *2P* versions but only a small amount from *ΔERRS* and none from *Δfurin* or *Tri*, both of which have deletions of the furin recognition site (Fig. 2A). Similarly, S2 was detected by anti-FLAG antibody in lysates from cells infected with *WT*, *D_614_*G and their *2P* versions but not from *Δfurin*- or *Tri* (Fig. 2B). Relatively small amounts of possibly aggregated higher molecular weight S was detected by anti-FLAG antibody in cells infected with *WT* and *D614G* (Fig. 2B).

**Fig. 2.**
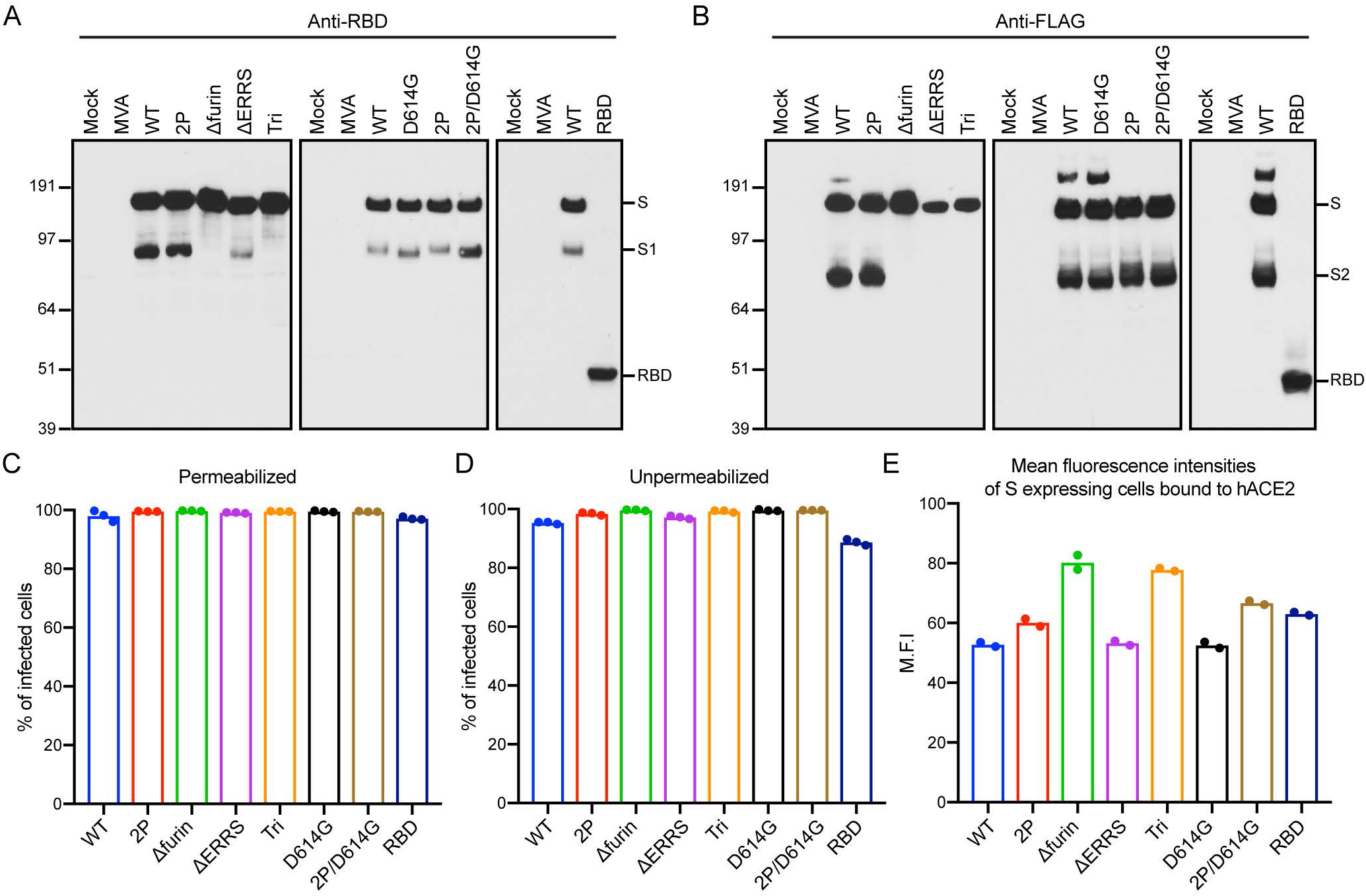
Expression of modified S proteins. (**A, B**) Western blots. HeLa cells were mock infected or infected with 5 PFU per cell of indicated rMVA for 18 h and total lysates were analyzed by SDS polyacrylamide gel electrophoresis. After membrane transfer, the proteins were detected with antibody to RBD or FLAG. The positions and masses in kDa of marker proteins are indicated on left and the positions of S, S1, S2 and RBD on right. (**C, D**) Flow cytometry. HeLa cells were infected in triplicate and permeabilized or stained directly with anti-SARS-CoV-2 Spike RBD mAb followed by APC-conjugated goat anti-mouse IgG. Infected cells were identified by GFP fluorescence and the percent that express S was determined by antibody staining. Bars represent the geometric mean. (**E**) Mean fluorescent intensities. HeLa cells were infected in duplicate and incubated with soluble hACE2 followed by Alexa Fluor 647-conjugated anti-hACE2 antibody. Cells that express S were identified as in the previous panels and the intensity of anti-hACE2 antibody determined. A representative of two experiments is shown.

Expression of the S proteins in HeLa cells that were infected with the rMVAs was also evaluated by flow cytometry. After permeabilization, virtually 100% of infected cells distinguished by GFP fluorescence were stained by a mouse anti-RBD mAb (Fig. 2C). In the absence of permeabilization, nearly 100% of the cells expressing full length S and nearly 90% expressing the RBD were stained indicating cell surface expression (Fig. 2D). Control experiments with unmodified parental MVA demonstrated no significant staining with or without permeabilization.

Human angiotensin converting enzyme (hACE2) is a cell receptor for CoV-2 (1, 37). The binding of soluble hACE2 to S proteins expressed on the surface of cells infected with rMVAs was analyzed as an indication of their appropriate folding. Binding of hACE2 to all constructs is shown in histograms (Fig. S1). The mean fluorescence intensities of S-expressing cells were similar except for the slightly higher value with *Δfurin* and *Tri* (Fig. 2E). We concluded that the WT and modified S proteins were all highly expressed on the surface of infected HeLa cells and potentially capable of eliciting immune responses.

### Binding and neutralizing antibodies induced by rMVAs

To compare their immunogenicity, each rMVA was inoculated intramuscularly (IM) into BALB/c mice at 0 time and again at 3 weeks. Some mice received purified RBD protein in QS21 adjuvant for priming and boosting or as a boost for mice primed with an rMVA. Binding antibody was measured by ELISA, using wells coated with purified 2P-stabilized S protein, at 3 weeks after the prime and 2 weeks after the boost. Binding antibodies were detected after the first immunization in all cases and increased by more than 1 log following a boost with the same vector (Fig. 3A, B). The ELISA titers for immunizations with all rMVAs were similar. Lesser binding, representing cross-reactivity, was obtained with sera from mice immunized with rMVA expressing the CoV-1 S (Fig. 3A) (19) and no binding above the base line was detected with sera from mice immunized with the parental MVA (Fig. 3A, B). Sera from mice immunized with the RBD protein and adjuvant exhibited low or no binding to S after the prime and more than a log less binding than any of the rMVAs expressing S after boosting with the same protein (Fig. 3A). Nevertheless, the RBD protein effectively boosted mice primed with rMVAs (Fig. 3A, B). The inability of RBD protein to induce binding antibody in naive mice was probably due to low immunogenicity of a soluble protein rather than to the truncation of the S sequence since the titer obtained after one vaccination with *RBD,* which encodes a membrane bound version, was similar to titers obtained with full-length S (Fig. 3B).

**Fig. 3.**
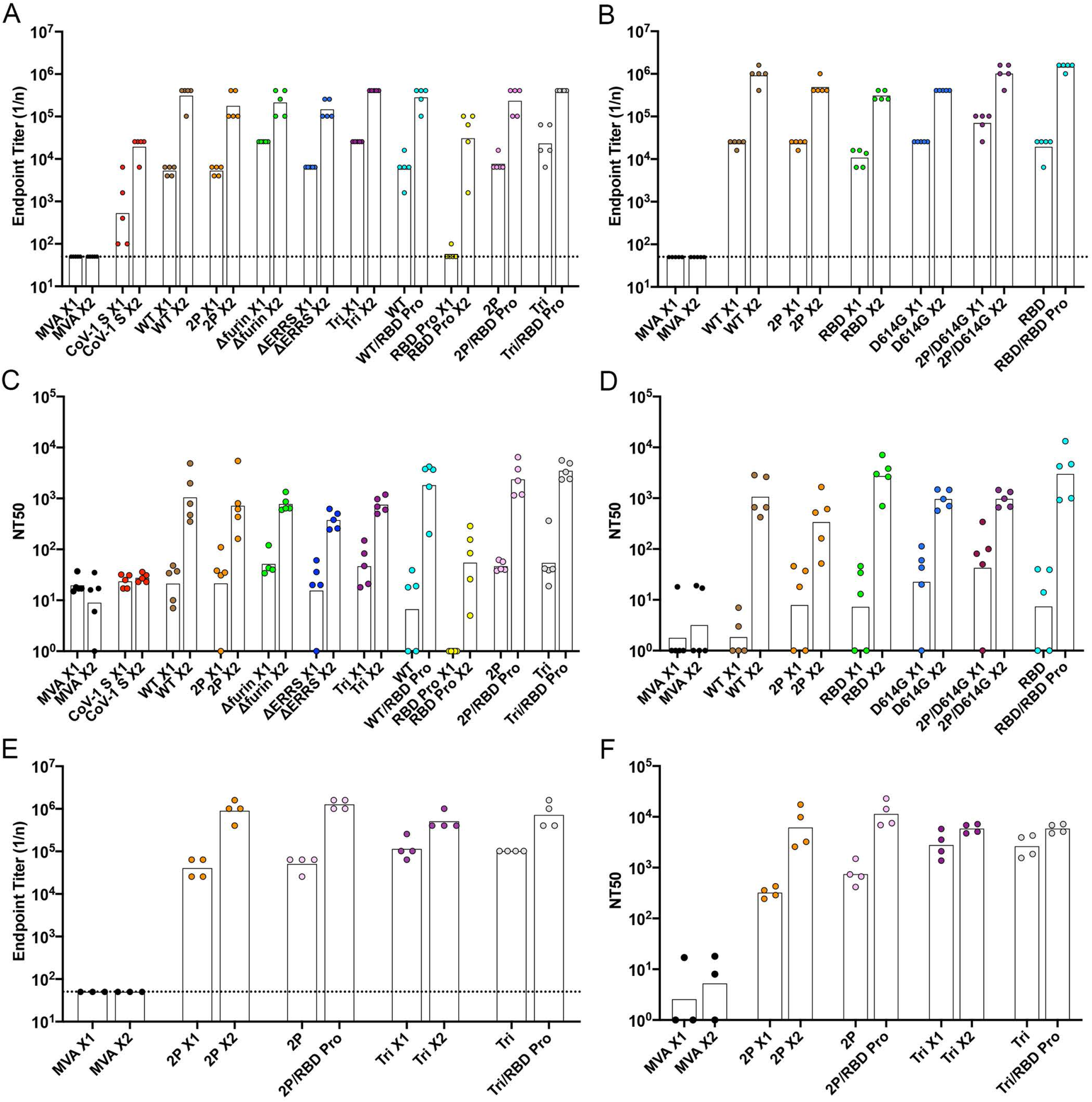
Binding and neutralizing antibody responses. BALB/c (**A-D**) or C57BL/6 (**E, F**) mice were primed and boosted 3 weeks later with 10^7^ PFU of parental MVA, rMVA expressing CoV-1 S or rMVA expressing CoV-2 WT S or modified S proteins in each hind leg or with 10 μg of RBD protein in QS21 adjuvant in the left hind leg. Mice were bled before vaccination, at 3 weeks (just before the boost), and at 2 weeks after the boost. Antibody binding to S was determined by ELISA. Reciprocal end point binding titers are shown in **A, B** and **E**. Dotted lines indicate limit of detection. Pseudovirus neutralization titers are shown in **C, D** and **F** and are plotted as NT50. In all panels, the tops of bars are the geometric mean titers. Abbreviations: X1 refers to sera collected 3 weeks after prime; X2 refers to sera collected two weeks after homologous boost; /indicates heterologous boost with RBD protein.

Neutralizing titers of the serum samples from BALB/c mice were determined using a lentiviral pseudotype assay (11, 38). Low or no neutralization was detected at 3 weeks after priming but increased markedly at 2 weeks after the rMVA boosts to mean levels of ~10^3^ NT50 (Fig. 3C, D). The RBD protein boosts elicited NT50 titers that were consistently higher than the rMVA boosts. Three samples of patient sera that had reference CoV-2 NT50 titers of 1280, 320, and 320 were found to have pseudovirus NT50 titers of 3209, 370 and 482, respectively. Thus, the rMVAs produced neutralizing antibody that was in the high range for patient sera.

We also determined binding and neutralization antibodies in sera from C57BL/6 mice that were immunized with *2P* and *Tri* and boosted with the same rMVAs or with RBD protein. Both the mean binding and neutralization titers were consistently higher in sera of C57BL/6 mice compared to BALB/c mice (Fig. 3E, F). The difference between the mouse strains was most significant for the neutralization titer after the primes (p<0.001). A time course indicated that binding and neutralizing antibody increased greatly between 1 and 3 weeks after the first immunization in C57BL/6 mice (Fig. S2). An additional experiment showed that soluble S proteins and RBD from a variety of sources boosted binding and neutralizing antibodies in C57BL/6 mice (Fig. S3).

High ratios of IgG2a and IgG2c to IgG1 in BALB/c and C57BL/6 mice, respectively, is indicative of a Th1 type anti-viral response (39). We determined the IgG subclasses of S-specific antibodies induced by an MVA-based vector and by RBD protein administered with QS21 adjuvant. The subclasses were determined by ELISA in which serum samples from vaccinated mice were added to 96-well plates that had immobilized full-length S. Following this, isotype specific secondary antibodies conjugated to horse radish peroxidase (HRP) were added. Table 1 shows that BALB/c and C57BL/6 mice that were primed and boosted by rMVA *2P* made IgG1, IgG2b, IgG2a or IgG2c and IgG3, but no detectable IgA antibody. The highest values were to IgG2a and IgG2c in BALB/c and C57BL/6, respectively (40, 41). The isotypes produced in the hACE2 mice, which were backcrossed to C57BL/6 and used in a later section of the paper, were similar to that of C57BL/6. However, the biggest difference was between the RBD protein prime and boost and immunizations with rMVA (Table 1). The protein only immunizations elicited a, predominance of IgG1 giving a clear Th2 response. Nevertheless, the RBD protein following rMVA *2P* boosted the Th1 response in both C57BL/6 and BALB/c mice.

**Table 1.**
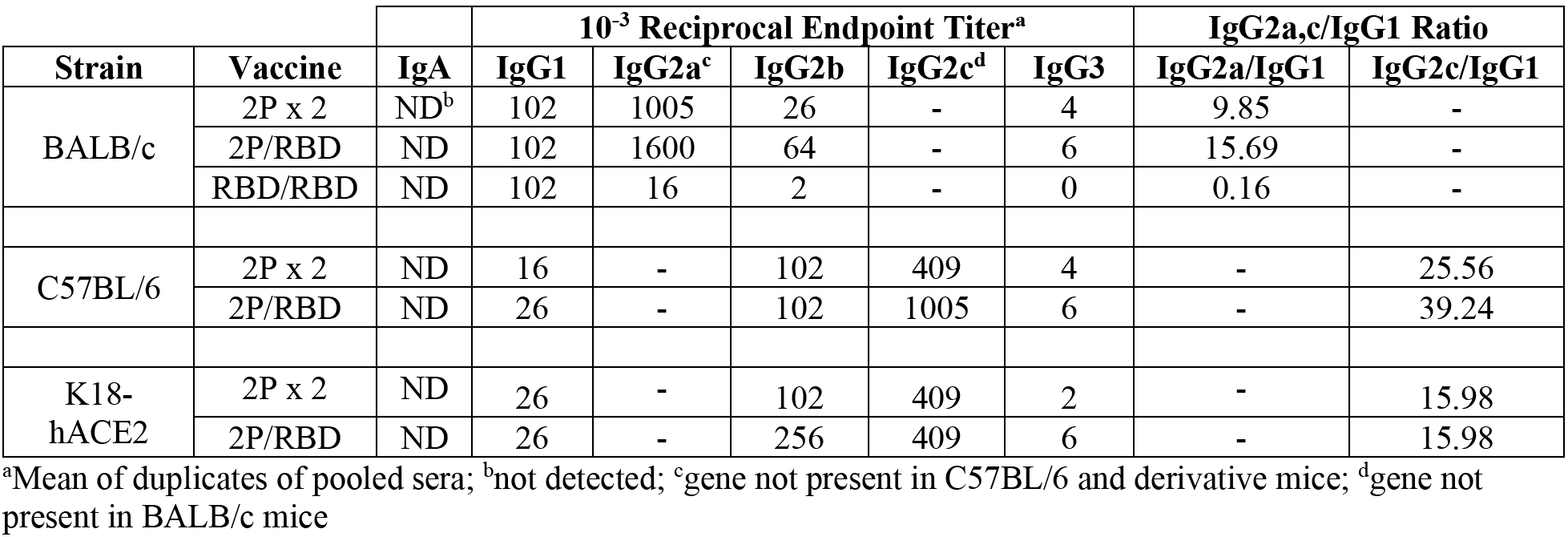
Isotype analysis of anti-S antibodies

| Strain | Vaccine | 10<sup>-3</sup> Reciprocal Endpoint Titer<sup>a</sup> |  |  |  |  |  | IgG2a,c/IgG1 Ratio |  |
| --- | --- | --- | --- | --- | --- | --- | --- | --- | --- |
|  |  | IgA | IgG1 | IgG2a<sup>c</sup> | IgG2b | IgG2c<sup>d</sup> | IgG3 | IgG2a/IgG1 | IgG2c/IgG1 |
| BALB/c | 2P x 2 | ND<sup>b</sup> | 102 | 1005 | 26 | - | 4 | 9.85 | - |
|  | 2P/RBD | ND | 102 | 1600 | 64 | - | 6 | 15.69 | - |
|  | RBD/RBD | ND | 102 | 16 | 2 | - | 0 | 0.16 | - |
| C57BL/6 | 2P x 2 | ND | 16 | - | 102 | 409 | 4 | - | 25.56 |
|  | 2P/RBD | ND | 26 | - | 102 | 1005 | 6 | - | 39.24 |
| K18-hACE2 | 2P x 2 | ND | 26 | - | 102 | 409 | 2 | - | 15.98 |
|  | 2P/RBD | ND | 26 | - | 256 | 409 | 6 | - | 15.98 |

<sup>a</sup>Mean of duplicates of pooled sera; <sup>b</sup>not detected; <sup>c</sup>gene not present in C57BL/6 and derivative mice; <sup>d</sup>gene not present in BALB/c mice

### Stimulation of Specific T cells

An *ex vivo* stimulation protocol was used to identify T cells specific for S following immunization. The sequences of an array of CoV-2 S peptides obtained from BEI Resources were compared to peptides that were previously found to be positive (42). As the peptides were not identical in the two libraries, we tested peptide pools for their ability to stimulate CD3+CD8+IFNγ+ and CD3+CD4+IFNγ+ T cells from spleens of BALB/c mice that had been immunized with parental MVA and rMVA *WT* expressing CoV-2 S. The two S peptide pools with highest specific activity were #4 and #7, which contained peptides from the NTD and RBD portions of S1, respectively (Fig. 4A). None of the pools had high CD4+IFNγ+ specific activity for the rMVA expressing S. A similar screen was carried out with spleen cells derived from immunized C57BL/6 mice. Pool #7 was again most positive for CD8+INFγ+ T cells, whereas other pools showed less specific activity (Fig. 4B).

**Fig. 4.**
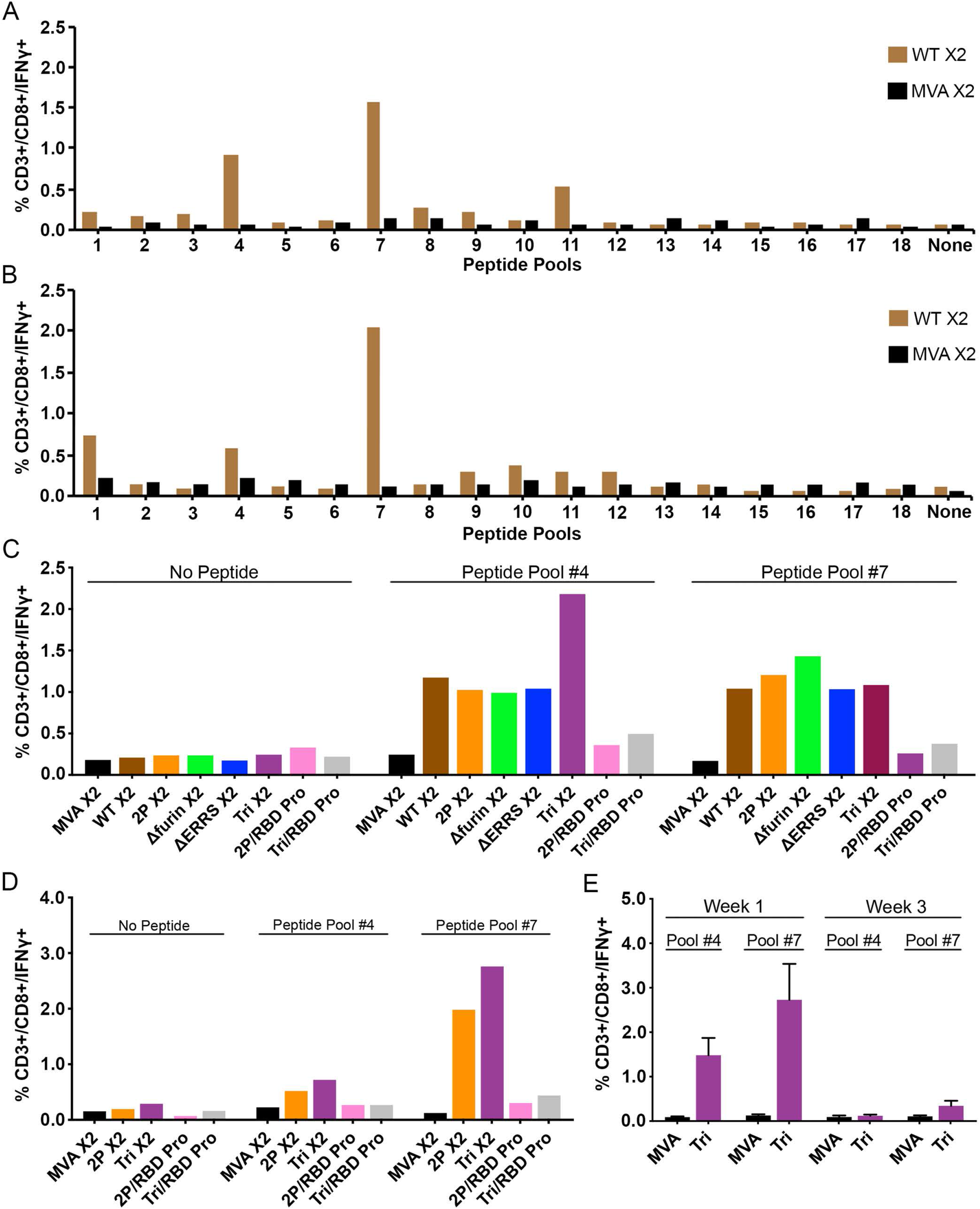
CD8+ T cell response. BALB/c mice **(A)** and C57BL/6 **(B)** mice were injected in each hind leg with 10^7^ PFU of unmodified MVA or rMVA expressing WT CoV-2 S at 0 time and again after 3 weeks. At 2 weeks after the boost, spleen cells were combined from 3-5 mice and stimulated with pools of peptides derived from CoV-2 S protein and treated with Brefeldin A. Cells were then stained for cell surface markers with mouse anti-CD3-FITC, anti-CD4-PE, and anti-CD8-PerCP. Cells were subsequently stained intracellularly with mouse anti-IFN--APC. CD3+CD8+IFNγ+ cells were enumerated by flow cytometry. BALB/c **(C)** and C57BL/6 **(D)** mice were primed with the indicated parental MVA or rMVA and boosted with the homologous rMVA or with RBD protein (Pro). Splenocytes from 4-5 mice were combined and stimulated with pool #4 and pool #7 peptides and then analyzed as in panels A and B. **(E)** C57BL/6 mice were primed with parental MVA or rMVA *Tri*. After 1 and 3 weeks the splenocytes of individual mice (n=4) were analyzed as in panel A. Abbreviations: X2 refers to splenocytes collected after homologous boost; /indicates heterologous boost with RBD protein. Standard deviations shown.

Next, we compared the percentages of splenic CD8+IFNγ+ T cells following priming with several different rMVA S constructs followed by homologous rMVA or RBD protein boosts. Spleen cells from mice immunized with parental MVA lacking S sequences and spleen cells that were not stimulated with peptide served as negative controls. Following priming and homologous boosting, peptide pool #7 stimulated T cells from mice immunized with each of the rMVA S constructs (Fig. 4C). A similar result was obtained with peptides from pool #4. In comparison, the CD8+IFNγ+ T cell numbers following boosts with RBD protein were much lower after stimulation with pool #4 or #7 than after rMVA boosts (Fig. 4C). The same pattern occurred in C57BL/6 T cells: the homologous prime boosts with *2P* and *Tri* were higher than, with RBD protein boosts (Fig. 4D). To better understand the difference between the boosts with MVA vectors and protein, we determined the duration of the T cells response following priming with MVA vectors. A dramatic drop in CD8+IFNγ+ T cell numbers occurred between 1 and 3 weeks (Fig. 4E), indicating a requirement for boosting by an MVA vector to restore elevated T cells.

### Protection against Intranasal (IN) CoV-2 infection

Transgenic mice that express hACE2 regulated by the cytokeratin 18 (K18) gene promoter (K18-hACE2) (43) are highly susceptible to IN CoV-2 infection (44). High levels of virus are present in the lungs within a few days and severe weight loss occurs by day 5 or 6 with animals becoming moribund. In our experiments, hACE2 transgenic mice were immunized by IM inoculation with *2P* or *Tri* and boosted 3 weeks later with the homologous rMVA or with RBD protein in adjuvant (Fig. 5A). Control mice were unvaccinated (naive) or primed and boosted with the parental MVA lacking CoV-2 sequences. Binding antibody to full length S was detected 3 weeks after the *2P* and *Tri* primes and was boosted up to 10-fold with homologous or protein boosts (Fig. 5B). Isotype analysis indicated that the binding antibody was IgG2c > IgG2b > IgG1 > IgG3 and no detectable IgA (Table 1), indicating a strong Th1 response in the mice receiving *2P* or *Tri* and homologous or heterologous boosts. In contrast to binding antibody, CoV-2 neutralizing antibody was boosted by RBD protein but not appreciably by the rMVAs (Fig. 5C). To help explain the latter results, we compared the MVA neutralizing antibodies of hACE2 mice after priming and boosting with rMVA *2P* as well as after boosting with RBD protein. The *2P* prime elicited high MVA neutralizing antibody, which was increased by more than a log after the homologous boost but not by the RBP protein boost (Fig. S4). We suspect that the primary antibody response to the rMVA attenuated the subsequent MVA infection and that the boost in MVA neutralizing antibody and CoV-2 S binding antibody were due in part to the virus particles and associated membranes of the virus inoculum. Boosting with the RBD protein, however, was not attenuated by priming with the rMVA and stimulated a strong anamnestic response to CoV-2 S.

**Fig. 5.**
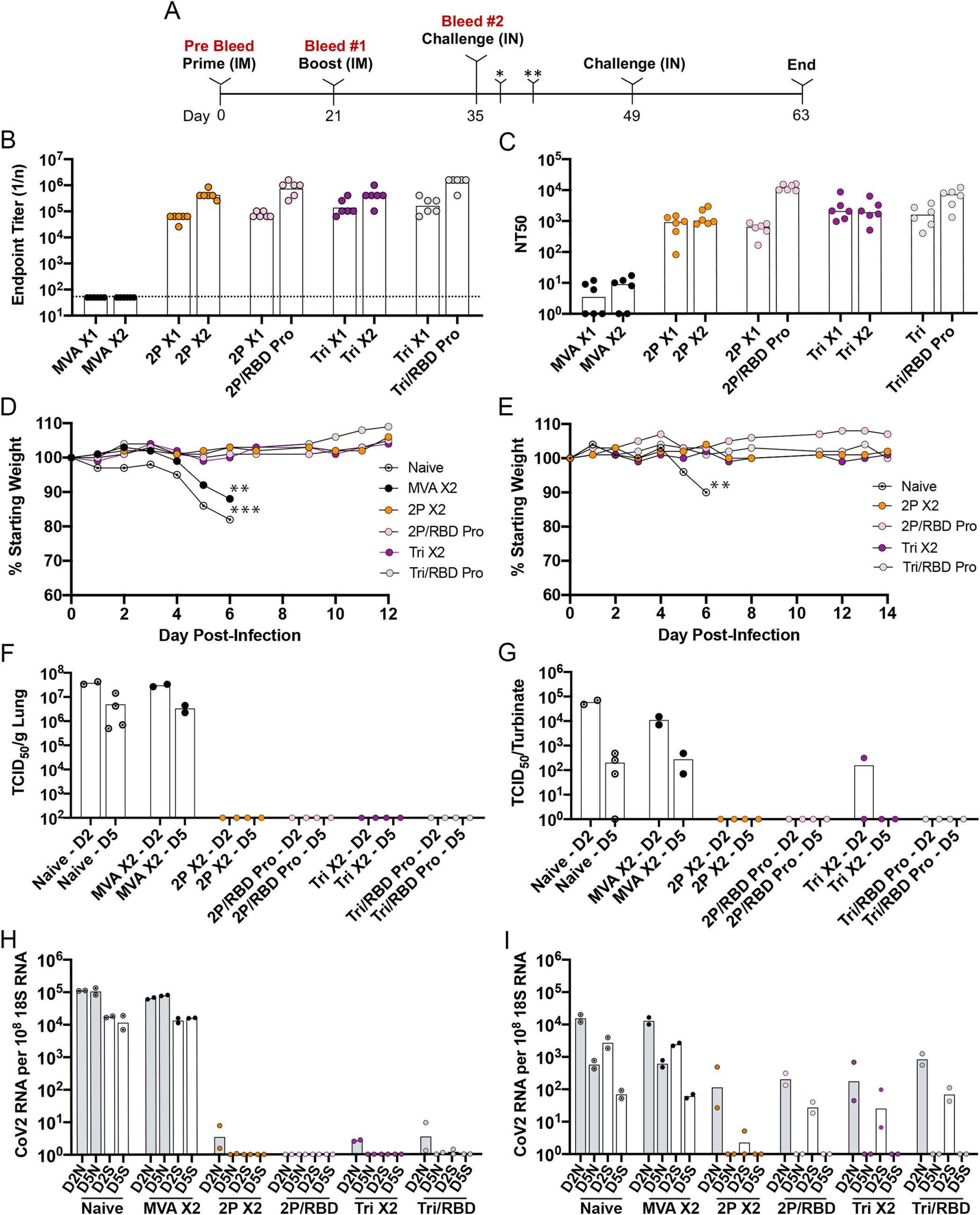
Challenge of transgenic mice following prime and boost vaccinations. (**A**) Protocol consisted of vaccinating five groups of six K18 hACE2 mice (female, 7 weeks old) IM in each hind leg with 10^7^ PFU of MVA or rMVA on days 0 (prime) and 21 (boost) and challenging unvaccinated (naive) and vaccinated mice with 10^5^ TCID50 of CoV-2 IN on day 35. A second IN challenge of surviving mice and two added naive mice was performed 2 weeks after the first challenge. Mice were weighed daily and observed for signs of morbidity. Mice were bled before vaccination (pre-bleed) and before the boost and 2 weeks after the boost. After challenge, 2 mice from each group were sacrificed on days 2 (*) and 5 (**) to determine the amounts of CoV-2 virus and subgenomic RNA. (**B**) Binding antibody was determined by ELISA on serum from each of the six mice of each vaccinated group and plotted as 1/end-point dilution. Dotted line indicates limit of detection. Abbreviations: X1 refers to sera collected 3 weeks after prime; X2 refers to sera collected 2 weeks after homologous boost; /indicates heterologous boost with RBD protein. (**C**) Neutralizing antibody was determined by a pseudovirus assay on serum from each of the six mice in each group and plotted as NT50. (**D**) Weights of surviving mice were determined daily and plotted as per cent of starting weight. Asterisks indicate number of mice that died or euthanized on a specific day. Data for one of the naive mice was obtained in a preliminary experiment. (**E)** Weights were determined following the second challenge of surviving mice and two naive mice. (**F**) Virus titers in lung homogenates obtained on days 2 and 5 were determined by end point dilution and plotted as TCID50 per gram of tissue. The lower two data points for naive mice on day 2 were determined in a preliminary experiment. (**G**) Virus titers in nasal turbinate homogenates obtained on days 2 and 5 were determined as in panel F and plotted as TCID50 per sample. (**H**) RNA was isolated from lung homogenates on days 2 and 5. CoV-2 N and S subgenomic RNAs were determined by ddPCR and plotted as copies per 10^8^ copies of 18s rRNA in the same sample. (**I**) RNA was isolated from nasal turbinate homogenates as described in panel H.

At 2 weeks after the boosts, the hACE2 mice were infected IN with 1×10^5^ TCID_50_ of CoV-2. The naive and parental MVA immunized mice lost weight by day 5 and were moribund on day 6 (Fig. 5D). The similarity between the two controls indicated that non-specific innate immune responses due to the parental MVA were not significantly protective at the time of challenge. In contrast, regardless of the boost, the rMVA vaccinated mice lost no weight and appeared healthy throughout the experiment. The surviving mice were re-challenged with CoV-2 after 2 weeks and again showed no weight loss, whereas additional naive mice succumbed to the virus infection by day 6 (Fig. 5E).

The lungs of naive and parental MVA immunized control mice had high titers of infectious CoV-2 on day 2, which dropped slightly on day 5, whereas no infectious virus was found in any of the 16 rMVA vaccinated mice at either time regardless of the prime or boost (Fig. 5F). The virus titers in the nasal turbinates of control mice peaked at day 2 but were still elevated at day 5 (Fig. 5G). In contrast, only one rMVA vaccinated mouse had a low level of virus in the turbinates that was more than 2 logs lower than controls on day 2 and none had detectable virus on day 5.

Subgenomic N and S mRNAs were analyzed by digital droplet PCR (ddPCR) using specific primers to distinguish newly synthesized RNA from input viral RNA. High levels of N and S mRNA were found in lungs of both control groups on days 2 and 5 (Fig. 5H). In contrast S mRNA was not detected in the lungs of rMVA vaccinated animals at either time and the more, abundant N mRNA was barely detected in a few animals at 4.5 to 5 logs lower than controls on day 2 and none on day 5 (Fig. 5H). N and S mRNAs were also detected in the nasal turbinates of control mice on both days but had decreased about a log between the two times (Fig. 5I). In rMVA vaccinated mice, N and S mRNAs were only detected on day 2 and the amounts were 43- and 85-fold lower, respectively, than the controls. Statistical significance (p<0.004) was determined by combining the values for N mRNAs on day 2 from both control groups and comparing that to the combined values from all rMVA vaccinated mice. The same p value was obtained when the S mRNAs were compared. Neither N nor S mRNA was detected in the nasal turbinates of any of the rMVA vaccinated mice on day 5. Thus, all rMVA vaccinated mice exhibited a high degree of protection against CoV-2 regardless of whether they were primed with *2P* or *Tri* and boosted with the homologous rMVA or RBD protein.

### Single vaccination

The similar levels of neutralizing antibody after priming and boosting with *2P* and *Tri*, led us to investigate whether a single rMVA vaccination would be sufficient to protect hACE2 mice against an intranasal challenge with CoV-2. The hACE2 mice were vaccinated with parental MVA or *Tri* and challenged 3 weeks later. Prior to challenge, the elevated binding and neutralizing antibody levels (Fig. 6A, B) were consistent with previous experiments. Following IN administration of CoV-2, mice that received the parental MVA suffered severe weight loss and became moribund, whereas the mice that received *Tri* remained healthy (Fig. 6C). Moreover, no infectious CoV-2 or subgenomic N or S mRNA was detected in lungs or nasal turbinates of the rMVA vaccinated mice on day 5 (Fig. 6D-F).

**Fig. 6.**
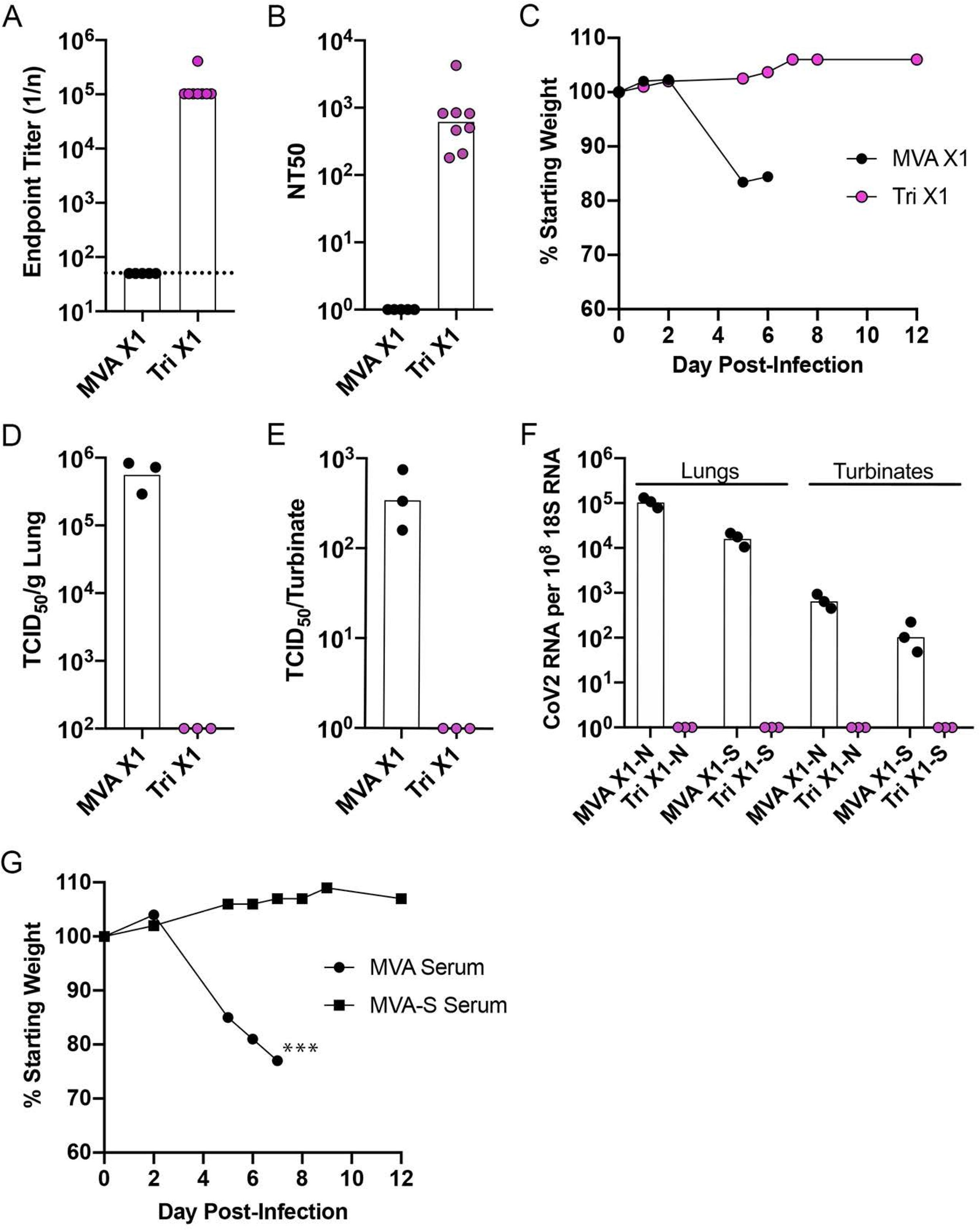
Challenge of transgenic mice after a single vaccination and after passive transfer of immune sera. (**A-F**) K18-hACE2 mice (female, 7 weeks old) were vaccinated IM on each hind leg with 10^7^ PFU of MVA (n=5) or rMVA *Tri* (n=8) and 5 mice of each group were challenged 3 weeks later with 10^5^ TCID50 of CoV-2 IN. (**A**) S-binding antibody prior to challenge. (**B**) Neutralizing antibody prior to challenge. (**C**) Weights of mice after challenge. (**D**) Lung virus titers. (**E**) Nasal turbinate virus titers. (**F**) N and S subgenomic RNA copies per 10^8^ copies of 18s RNA in lung and nasal turbinates. **(G)** Passive serum transfer. K18-hACE2 mice were injected IP with 0.4 ml of pooled serum from mice vaccinated with parental MVA (n=3) or with MVA-S (n=4) and challenged with 10^5^ TCID50 of CoV-2 IN. Mice were weighed on the indicated days and values plotted as percent of starting weight of each mouse. Asterisks indicate number of mice that died or were sacrificed due to morbidity on day 7.

### Protection of hACE2 mice by passive transfer of serum

A passive transfer experiment was carried out to determine whether antibody induced by vaccination with rMVA S vectors is sufficient to protect against lethal infection with CoV-2. Sera were pooled from mice that had been vaccinated by priming and boosting with parental MVA or rMVA expressing WT S. Aliquots were injected into the peritoneum of hACE2 mice, which were challenged with CoV-2 approximately 24 h later. A few hours before the challenge, the mice were bled and the pseudotype neutralizing titers of 160 NT50 were found for each of the mice that received the immune serum, well below the titers of mice that received rMVA vaccinations. Nevertheless, the mice showed no signs of weight loss or ill health following inoculation of CoV-2 (Fig. 6G).

## Discussion

The CoV-2 S protein is the major target of neutralizing antibodies. In an attempt to optimize the synthesis and immunogenicity of S, we constructed a panel of rMVAs that expressed unmodified S or S with one or multiple modifications. However, little or no difference was found in the cell surface expression of the variant forms of S and each of the rMVAs stimulated similar levels of antibody that bound S in an ELISA and neutralized a CoV-2 S pseudotype virus. A common feature of all the rMVAs was surface expression of the RBD as shown by interaction with an anti-RBD mAb and soluble hACE2.

Isotype analysis indicated that the rMVAs induced a well-balanced predominantly Th1 type response with IgG2a or IgG2c (depending on the mouse strain) > IgG2b > IgG1 > IgG3, which is the usual order following a viral infection and by IFNγ stimulation (45–47). IgG2a, IgG2b and IgG2c have similar functions and are able to fix complement and activate Fc receptors to promote virus clearance, whereas IgG1 may limit inflammation (47). Lower levels, of binding and neutralizing antibodies were detected following priming and boosting with purified soluble RBD protein in QS21 adjuvant. In addition, IgG1 was predominant when mice were immunized with RBD protein. Nevertheless, higher binding and neutralizing titers were obtained after priming with an rMVA and boosting with RBD protein rather than the homologous rMVA. In part, this may be explained by attenuation of rMVA boosts by immunity to the live virus vector that was generated during the prime. Extending the interval between the first and second rMVA vaccinations to allow the anti-MVA immunity to decline might enhance boosting. Interestingly, the predominance of IgG2a and IgG2c was maintained when RBD was used as the boost for rMVAs, suggesting that the protein stimulated an anamnestic response. The trade-off, however, was a lower CD8+IFNγ+ T cell response with the heterologous protocol.

The neutralizing antibody titers obtained with the rMVAs (NT50 of ~10^3^ or higher with the RBD protein boost) compared favorably with that achieved by mRNA immunizations. Using the same lentivirus pseudovirus protocol and reagents, the neutralizing titers for immunizations after a prime and boost with mRNA encoding the 2P-modified S reached 819, 89 and 1115 reciprocal IC50 geometric mean titer for BALB/c, C57BL/6 and B6C3F1/J mice, respectively (11) and pseudovirus neutralizing titers of ~340 NT50 were obtained with 100 to 250 μg of mRNA in a phase 1 clinical study (48). How well rMVA S constructs will do in other animal models and humans remains to be determined.

K18-hACE2 mice were chosen for CoV-2 protection studies because of their susceptibility to severe disease including lung inflammation and death (43, 44). Studies were performed in which the mice were challenged after priming and boosting or after just priming. For the prime-boost study, the mice were first vaccinated with rMVAs *2P* or *Tri* and boosted with the homologous rMVA or with RBD protein in QS21 adjuvant. The control naive mice and mice immunized with the parental MVA lost weight and exhibited signs of morbidity within 5 to 6 days after IN inoculation with CoV-2, whereas the challenged mice of all rMVA vaccination groups remained healthy and without weight loss. The rMVA vaccinated mice also resisted a second challenge two weeks later. High virus titers were present in the lungs of the control mice on day 2 and slightly lower on day 5, whereas no virus was detected in the lungs of any of the vaccinated mice. Although CoV-2 was isolated from the nasal turbinates of all control mice on days 2 and 5, only one of eight rMVA vaccinated mice had a low amount of virus on day 2 and none of eight had detectable virus on day 5. High levels of subgenomic N and slightly lower S mRNAs were detected in the lungs of control mice, whereas only traces of N and no S were found in a minority of rMVA vaccinated mice on day 2 only. However, all rMVA vaccinated mice had low levels of subgenomic RNA in the nasal turbinates on day 2 compared to control mice, which was cleared by day 5. Mice challenged after a single rMVA immunization were also protected and had no detectable virus or subgenomic RNAs in the lungs or nasal turbinates on day 5. The detection of low amounts of RNA in the nasal turbinates that was subsequently cleared indicated that sterilizing immunity had not been obtained by systemic rMVA vaccination and would likely require local immunization.

Since the MVA vectors stimulated both antibody and T cells, a passive immunization experiment was carried out to evaluate the protective role of antibody alone. Serum from BALB/c mice that had been vaccinated with MVA expressing WT S was injected intraperitoneally into K18-hACE2 mice resulting in NT50s of 160 prior to challenge. Following challenge, the mice remained healthy and lost no weight indicating that antibody is sufficient for protection in this model and that the level of neutralizing antibody elicited by active immunization is considerably higher than necessary.

In the present study, we evaluated rMVA CoV-2 vaccines administered IM with homologous or protein boost protocols. Previous studies have shown enhanced responses when rMVAs were combined with recombinant DNA or other virus vectors (49–51). For example, a filovirus vaccine consisting of an Ad26 vector followed by a rMVA was safe and immunogenic in a phase 2 trial (51). The stability of both Ad26 and rMVA compared to mRNA vaccines, which must be kept frozen except for short periods, is an advantage for global distribution. The rMVA component was shown to remain stable for 24 months frozen, 12 months at 2-8°C and up to 6 h at 40°C in a syringe needle (52). Another point to consider is the route of administration, which can affect the ability of vaccines to prevent spread. In addition to IM and subcutaneous routes, rMVAs can be administered orally, IN and by aerosol (53–60). Although IM administration of the rMVA CoV-2 S vectors greatly reduced and rapidly eliminated virus replication in the nasal turbinates, it will be of interest to determine whether IN administration would prevent replication entirely.

## Materials and Methods

### Cells

Cells were maintained at 37°C in 5% CO_2_ humidified incubators. HeLa cells (ATCC CCL-2) and Vero E6 cells (ATCC CRL-1586) were grown in Dulbecco’s modified eagle’s medium (DMEM) supplemented with 8% fetal bovine serum (FBS, Sigma-Aldrich), 2 mM L-glutamine, 100 U/ml of penicillin, and 100 μg/ml of streptomycin (Quality Biological). 293T-hACE2.MF cells were propagated in the above medium supplemented with 3 μg/ml of puromycin. Primary CEF prepared from 10-day old fertile eggs (Charles River) were grown in minimum essential medium with Earle’s balanced salts (EMEM) supplemented with 10% FBS, 2 mM L-glutamine, 100 U/ml of penicillin, and 100 μg/ml of streptomycin.

### Mice

Five-to six-week-old female BALB/cAnNTac and C57BL/6ANTac were obtained from Taconic Biosciences and B6.Cg-Tg(K18-ACE2)2Prlmn/J from Jackson Laboratories. Mice were separated into groups of 2-5 animals in small, ventilated microisolator cages in an ABSL-2 facility and used after 1-5 additional 5 weeks.

### Construction of Recombinant Viruses

DNA encoding the CoV-2 S protein (QHU36824.1), with a C-terminal 3xFLAG tag and modified by removing four VACV early transcription termination signals (TTTTTNT) and runs of four or more consecutive Gs or Cs, was chemically synthesized (Thermo Fisher). Another construct (*Tri*) with the proline substitutions (K986P/V987P), furin recognition site substitutions (aa 682–685 RRAR to GSAS), and C-terminal 19 aa deletion of ERRS was also synthesized. Constructs with these individual mutations were generated using Q5 Site-Directed Mutagenesis Kit (New England Biolabs). A 2-step PCR protocol using the Q5 mutagenesis kit was used to join nucleotides 1-117, 955-1782 and 3586-3819 to form the ORF for *RBD*. The DNAs were inserted into the pLW44 transfer vector (36) at the *Xma*I and *Sal*I sites, which placed the ORF under the control of the VACV modified H5 early late promoter and adjacent to the separate gene encoding enhanced GFP regulated by the VACV P11 late promoter.

To produce rMVAs, linearized plasmids were transfected into cells infected with MVA allowing recombination into the existing deletion III site in the MVA genome (36). The rMVAs were clonally purified by four successive rounds of fluorescent plaque isolation, propagated in CEF, and purified by sedimentation twice through a 36% sucrose cushion. The genetic purities of the recombinant viruses were confirmed by PCR amplification and sequencing of the modified, region. Titers of MVAs were determined in CEF by staining plaques with anti-VACV rabbit antibodies (36).

### Western Blotting

HeLa cells were infected with 5 PFU per cell of rMVAs for 18 h, washed once with phosphate buffered saline (PBS), then lysed in LDS sample buffer with reducing agent (Thermo Fisher). The lysates were dispersed in a sonicator for four 30 s periods; the proteins were resolved on 4 to 12% NuPAGE Bis-Tris gels (Thermo Fisher) and transferred to a nitrocellulose membrane with an iBlot2 system (Thermo Fisher). The membrane was blocked with 5% nonfat milk in Tris-buffered saline (TBS) for 1 h, washed with TBS with 0.1% Tween 20 (TBST), and then incubated at 4°C overnight with rabbit anti-CoV-2 RBD polyclonal antibody (Cat# 40592-T62, Sino Biological) or anti-FLAG M2 peroxidase antibody (Cat# A8592, MilliporeSigma) in 5% nonfat milk in TBST. The membrane that had been incubated with anti-RBD antibody was then incubated for 1 h with secondary antibody conjugated to horseradish peroxidase (Jackson ImmunoResearch). After washing, the membrane bound proteins were detected with SuperSignal West Dura substrate (Thermo Fisher).

### Detection of S Protein by Flow Cytometry

HeLa cells were infected with 5 PFU per cell of rMVAs. After 24 h, the infected cells were stained for intracellular and surface S in parallel. For intracellular staining, cells were fixed with Cytofix/Cytoperm (BD Biosciences), permeabilized with Perm/Wash Buffer (BD Biosciences), and incubated with anti-CoV-2 Spike RBD mAb (SARS2-02) (13) followed by APC-conjugated goat anti-mouse IgG antibody (Cat# 405308, BioLegend). For surface detection, the cells were stained directly using the same primary and secondary antibodies. The binding of hACE2 to surface expressed S protein on infected HeLa cells was detected by incubating with 100 ng/10^6^ cells of biotinylated human ACE2 protein (Cat# 10108-H08H-B, Sino Biological) followed by Alexa Fluor 647-conjugated anti-hACE2 antibody (Cat# FAB9332R, R&D Systems). The stained cells were acquired on a FACSCalibur cytometer using Cell Quest software and analyzed with FlowJo (BD Biosciences).

### Detection of S-Binding Antibodies by ELISA

CoV-2 S protein produced in HEK293 cells was obtained from the NIAID Vaccine Research Center or Sino Biological and diluted in cold PBS to a concentration of 1 μg/ml. Diluted S protein (100 μl) was added to each well of a MaxiSorp 96-well flat-bottom plate (Thermo Fisher). After incubation for 16-18 h at 4°C, the wells were washed 3 times with 250 μl of PBS + 0.05% Tween 20 (PBS-T, Accurate Chemical) and plates were blocked with 200 μl PBS-T + 5% Nonfat Dry Milk for 2 h at room temperature. During the blocking phase, a series of eight 4-fold dilutions of each mouse serum sample was prepared in blocking buffer. After blocking, plates were washed 3 times with 250 μl of PBS-T and 100 μl of each 4-fold dilution of serum was added to the appropriate well(s) and incubated for 1 h at room temperature. After incubation with serum, plates were washed 3 times with 250 μl of PBS-T. HRP-conjugated goat anti-mouse IgG (H+L) (Thermo Fisher) was diluted 1:4000 in blocking buffer and 100 μl of the secondary antibody was added to each well for 1 h at room temperature. For detection of antibody isotypes, peroxidase-conjugated isotype-specific antibodies were used (Thermo Fisher). Plates were washed 3 times with 250 μl of PBS-T and then 100 μl of KPL SureBlue TMB 1-component microwell peroxidase substrate (Seracare) was added to each well. The chemiluminescence reaction was stopped after 10 min by addition of 100 μl of 1N sulfuric acid. Spectrophotometric measurements were made at A_450_ and A_650_ using a Spectramax Plus 384 plate reader with Softmax Pro analysis software (Molecular Devices). Final endpoint titers, (1/n) for each sample were determined as 4-fold above the average OD of those wells not containing primary antibody (OD 0.03-0.04).

### Stimulation and Staining of Lymphocytes

Splenocytes from individual mice or pooled from 3-5 mice were suspended at 1.5×10^7^ cells/ml in RPMI (Quality Biological) supplemented with 10% heat-inactivated FBS, 10 U/ml penicillin, 10 μg/ml streptomycin, 2 mM L-glutamine, and 2 mM HEPES as previously described (61). Splenocytes (100 μl) were mixed with 100 μl of individual peptide pools in 96-well plates and incubated at 37°C for 1 h after which brefeldin A (Sigma Aldrich) was added and incubation continued for 4-5 h. Staining of cells was performed at 4°C. Fc receptors were blocked with anti-CD16/32 (Clone 2.4G2, a gift from Jack Bennink, NIAID) for 30 min. Surface staining was performed with anti-mouse CD3-FITC (Clone17A2; BioLegend), anti-mouse CD4-PE (Clone H129.19; BD Biosciences), and anti-CD8-PerCP-Cy5.5 (BD Biosciences) for 1 h. Cells were then fixed and permeabilized with Cytofix/Cytoperm solution and stained with IFNγ-APC (BD Biosciences) for 1 h. Cells were washed with PBS and suspended in PBS containing 2% paraformaldehyde. Approximately 100,000 events were acquired on a FACSCaliber cytometer using Cell Quest software and analyzed with FlowJo (BD Biosciences).

A peptide array was obtained from BEI Resources (catalog # NR-52402; SARS-Related Coronavirus 2 Spike (S) Glycoprotein). Each peptide was dissolved in DMSO at 10 μg/ml. A total of 18 peptide pools were prepared, containing 3-11 peptides per pool. For splenocyte stimulation, the final concentration of each peptide was 2 μg/ml. Peptides in the 2 positive pools were: Pool #4 (BEI peptides 32-41); Pool #7 (BEI peptides 61, 64, 77).

### Pseudovirus Neutralization Assay

The CoV-2 lentivirus pseudotype assay was carried out as described by Corbett et al. (11) using cells and plasmids obtained from the NIAID Vaccine Research Center. To determine neutralization titers, serum samples were heat inactivated for 30 min at 56°C and clarified by high speed microcentrifugation. The day before titration, 5,000 293T-hACE2.MF cells were seeded per well in 96-well white walled clear bottom tissue culture plates (Corning) in DMEM supplemented with 10% heat inactivated FBS, 2 mM L-glutamine, 100 U/ml penicillin, 100 μg/ml streptomycin) with 3 μg/ml of puromycin. For each serum sample, duplicate 4-fold dilution series were prepared in 96-well U-bottom plates (Corning) in DMEM supplemented with 5% heat inactivated FBS with the starting dilution being 1:20 in a final volume of 45 μl per well. The pseudovirus was thawed at 37°C and 45 μl of a dilution previously shown to exhibit a 1000-fold difference in luciferase between uninfected and infected cells was added to all wells except for controls. After 45 min at 37°C, the medium was aspirated and 50 μl sample-virus mixture was added to each well and incubated for 2 h at 37°C. DMEM (150 μl) supplemented with 5% heat inactivated FBS was added per well and plates were incubated for 72 h at 37°C. Medium was removed from the wells and the cells were lysed with 25 μl per well of 1X cell lysis reagent (Promega), shaken at 400 rpm for 15 min at room temperature. Luciferase reagent (50 μl, Promega) was added per well and 90 s later relative luciferase units (RLU) were read on the luminometer (EnSight, Perkin Elmer, 570 nm wavelength, 0.1 mm distance, 0.3 s read). NT50 were calculated using Prism (GraphPad Software) to plot dose-response curves, normalized using the average of the no virus wells as 100% neutralization, and the average of the no serum wells as 0%.

### MVA Neutralization Assay

A semi-automated flow cytometric assay was carried out as previously described (62) except for substitution of MVA expressing GFP for the WR strain of VACV. Briefly, ten 2-fold serial dilutions of heat-inactivated serum from vaccinated mice were prepared in a 96-well plate and 6.25×10^3^ PFU of MVA-GFP was added to each well and incubated at 37°C for 1 h. Approximately 10^5^ HeLa suspension cells were added to each well in the presence of 44 μg/ml of cytosine arabinoside. After 18 h at 37°C, the cells were fixed in 2% paraformaldehyde and acquired with a FACSCalibur cytometer using Cell Quest software and analyzed with FlowJo. The dilution of mouse serum that reduced the percentage of GFP-expressing cells by 50% (IC_50_) was determined by nonlinear regression using Prism.

### SARS CoV-2 Challenge Virus

SARS CoV-2 USA-WA1/2019 was obtained from BEI resources (Ref# NR-52281) and propagated in a BSL-3 laboratory using Vero E6 cells cultured in DMEM+Glutamax supplemented with 2% heat-inactivated FBS and penicillin, streptomycin, and fungizone by Bernard Lafont of the NIAID SARS Virology Core laboratory. The TCID50 of the clarified culture medium was determined on Vero E6 cells after staining with crystal violet and scored by the Reed-Muench method.

### Vaccination and Challenge Experiments

Prior to vaccination, the virus was thawed, sonicated twice for 30 s on ice and diluted to 2×10^8^ PFU/ml in PBS supplemented with 0.05% bovine serum albumin. In an ABSL-2 laboratory, 50 μl of diluted virus was injected IM into each hind leg of the animal for a total dose of 2×10^7^ PFU. Unless otherwise stated, baculovirus RBD protein provided by Eugene Valkov (NCI) was diluted to 0.2 mg/ml in PBS containing 0.3 mg/ml of QS-21 adjuvant (Desert King International, San Diego, CA) and 10 μg of RBD was, injected IM into the left hind leg. All mice scheduled to be infected with SARS-CoV-2 were transferred to an ABSL-3 laboratory a few days prior to virus challenge. The challenge stock of SARS-CoV-2 USA-WA1/2019 was diluted to 2×10^6^ TCID50/ml in PBS. Mice were lightly sedated with isoflourine and inoculated IN with 50 μl of SARS-CoV-2. After infection, morbidity/mortality status and weights were assessed and recorded daily for 14 days by the NIAID Comparative Medical Branch.

### Determination of CoV-2 in Lungs and Nasal Turbinates

At 2- and 5-days post-infection with CoV-2, lung and nasal turbinates were removed and placed in 1.5-2 ml of ice-cold Dulbecco’s PBS and weights of lungs were recorded. Tissues were homogenized for three 25 s intervals in ice water using a GLH-1 grinder equipped with a disposable probe and aerosol proof cap (Omni International). Homogenates were cleared of debris by centrifugation at 4,000 ×g for 10 min and the supernatants were transferred to sterile tubes and stored at −80°C. Clarified homogenates were thawed and titrated in quadruplicate on Vero E6 cells using 10-fold serial dilutions in 96-well microtiter plates. After 72-96 h, the plates were stained with crystal violet and scored using the Reed-Muench method to determine TCID50.

### Determination of CoV-2 RNA in Lungs and Nasal Turbinates

Immediately after homogenization of lungs and turbinates, 0.125 ml was transferred to sterile tubes, 0.9 ml Trizol (Thermo Fisher) was added and the mixture frozen. After thawing, RNA was extracted using the Trizol Plus RNA Purification Kit with Phasemaker tubes (Thermo Fisher) following the manufacturer’s instructions. Contaminating DNA was removed from the eluted RNA using the Turbo DNA-free kit (Thermo Fisher) and RNA was reverse-transcribed using the iScript cDNA, synthesis kit (Bio-Rad, Hercules, CA). CoV-2 S and N transcripts and 18s rRNA were quantified by ddPCR with specific primers (CoV-2 RNA Leader, Forward – CGA TCT CTT GTA GAT CTG TTC TCT AAA C; CoV-2 S, Reverse – TCT TAG TAC CAT TGG TCC CAG AGA; CoV-2 N, Reverse – GGT CTT CCT TGC CAT GTT GAG T; 18S, Forward – GGC CCT GTA ATT GGA ATG AGT C; 18S, Reverse – CCA AGA TCC AAC TAC GAG CTT) using an automated droplet generator and QX200 Droplet Reader (Bio-Rad). The values for CoV-2 S transcripts were normalized using the 18s RNA in the same sample.

### Passive Serum Transfer

Serum for passive transfer was obtained from 20 BALB/c mice that were inoculated IM with rMVA S (*WT*) and 10 BALB/c mice with parental MVA at 0 and 3 weeks. Two weeks after the boosts, the MVA S and control MVA sera were pooled separately. Four naive K18-hACE2 mice each received 0.4 ml of MVA S serum and three received 0.4 ml of the control MVA serum. The following day, mice were bled to determine levels of SARS-CoV-2 binding and neutralizing antibody. Approximately 4 h later, the mice were challenged IN with 10^5^ TCID_50_ of CoV-2. Mice were observed and weighed over the next two weeks.

### Safety and Ethics

All experiments and procedures involving mice were approved under protocol LVD29E by the NIAID Animal Care and Use Committee according to standards set forth in the NIH guidelines, Ansaimal Welfare Act, and US Federal Law. Euthanasia was carried out using carbon dioxide inhalation in accordance with the American Veterinary Medical Association Guidelines for Euthanasia of Animals (2013 Report of the AVMA Panel of Euthanasia). Experiments with SARS-CoV-2 were carried out under BSL-3 containment.

## Data Availability

Materials and data are available upon request.

## Acknowledgements

We thank Kizzmekia Corbett of the NIAID Vaccine Research Center for reagents and protocols, Eugene Valkov of the NCI for RBD protein, Bernard Lafont of NIAID for CoV-2 and use of the BSL-3 facility, Michael Holbrook of NIAID for patient serum, and Michael Diamond and Laura VanBlargan of Washington University in St. Louis for mouse mAbs. The technical staff of the NIAID Comparative Medical Branch provided animal care. The work was funded by the Division of Intramural Research, NIAID.

