## Supplemental Figures 1-4 for "MVA Vector Vaccines Inhibit SARS CoV-2 Replication in Upper and Lower Respiratory Tracts of Transgenic Mice and Prevent Lethal Disease"

### Supplemental Figure Legends

**Fig. S1.** Binding of hACE2. HeLa cells were infected with 5 PFU of rMVAs expressing WT or modified versions of S. After 24 h, the cells were incubated with soluble hACE2 and stained with Alexa Fluor 647-conjugated goat anti-hACE2 antibody. For the control, the addition of hACE2 protein was omitted. Histograms of two replicates are superimposed and M.F.I values of all infected cells are indicated.

**Fig. S2.** Time course of antibody production. C57BL/6 mice were vaccinated with MVA or rMVA *Tri*. Serum was collected before vaccination (Prebleeds) and 1 and 3 weeks after vaccination. **(A)** S-binding antibody determined by ELISA. **(B)** Neutralizing antibody determined by pseudovirus assay.

**Fig. S3.** Comparison of protein boosts. C57BL/6 mice were primed by IM injection with $10^7$ PFU of rMVA *2P* into each hind leg. After 3 weeks, the mice were boosted by IM injection with 10 μg of RBD produced in human cells (h-RBD, Genscript), baculovirus produced RBD (RBD#1, Sino Biological), baculovirus produced RBD (RBD#2, provided by Eugene Valkov, NCI), or soluble S protein produced in human cells (h-S, Sino Biological). Each protein was administered with 15 μg of QS21 adjuvant. The mice were bled after 2 weeks and binding antibody and neutralization titers determined by ELISA **(A)** or pseudovirus assay **(B)**, respectively.

**Fig. S4.** MVA neutralizing antibody. Sera obtained from individual hACE2 mice that were primed with MVA *2P* (2PX1) and boosted with MVA *2P* (2PX2) or with RBD protein (2P/RBD

Pro) were tested for the ability to neutralize MVA using a flow cytometry assay. The dilutions of mouse sera that reduced the percentage of GFP-expressing cells by 50% (IC~~50~~) were plotted.

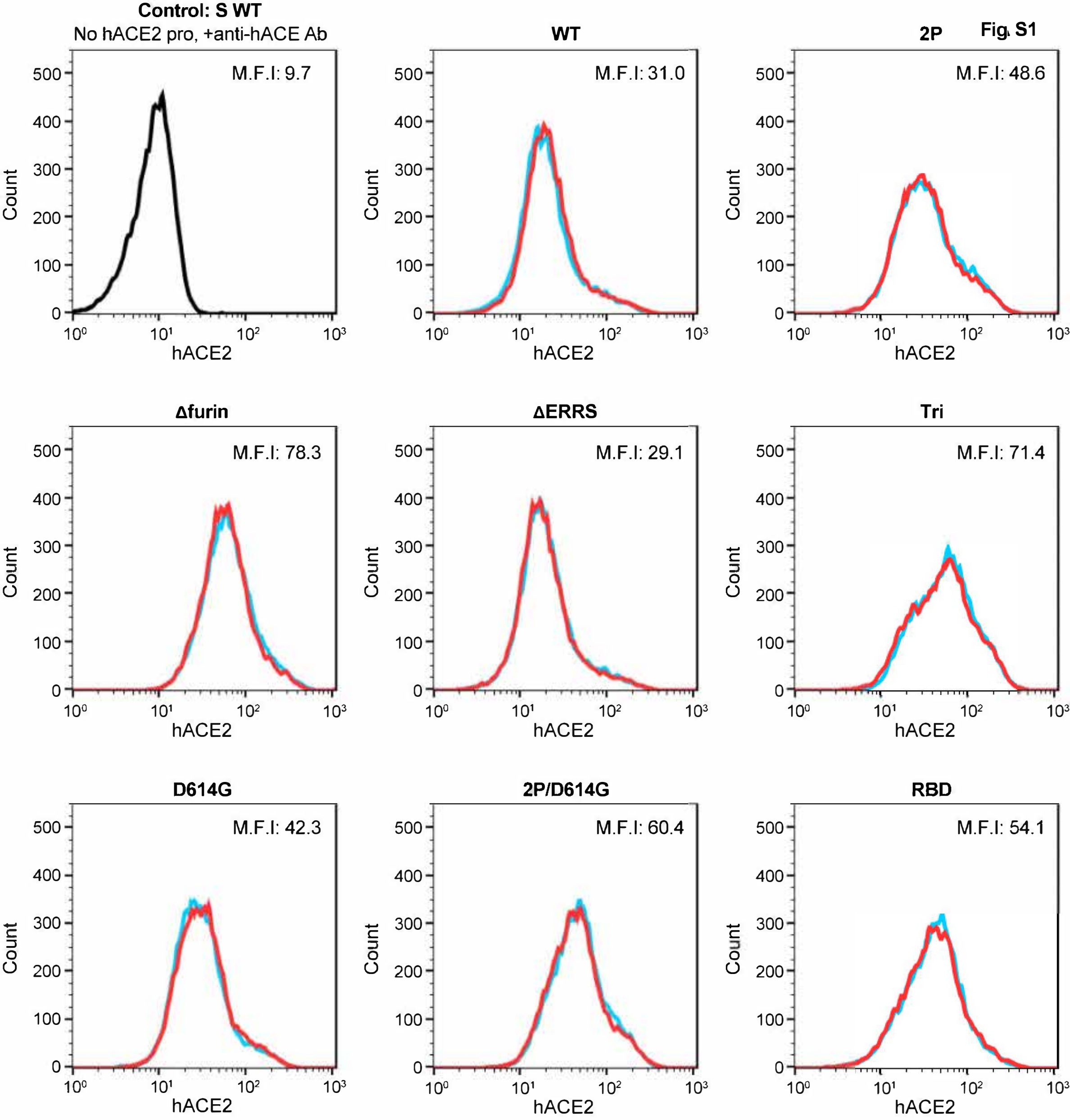
Fig. S1
Control: S WT
No hACE2 pro, +anti-hACE Ab
M.F.I: 9.7
WT
M.F.I: 31.0
2P
M.F.I: 48.6
Δfurin
M.F.I: 78.3
ΔERRS
M.F.I: 29.1
Tri
M.F.I: 71.4
D614G
M.F.I: 42.3
2P/D614G
M.F.I: 60.4
RBD
M.F.I: 54.1
Count
hACE2

Fig. S2

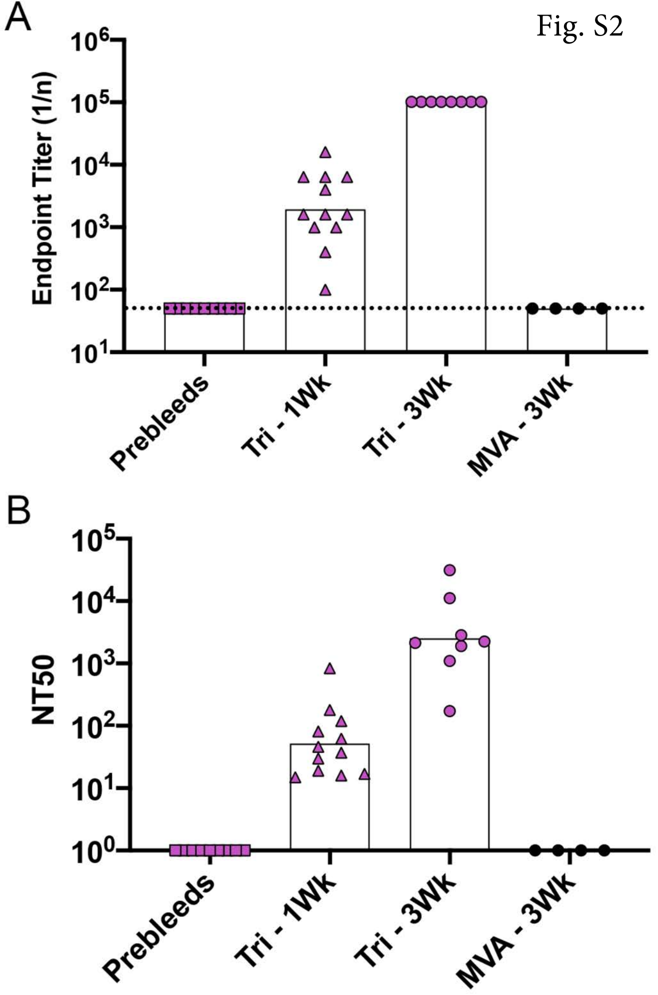

A
Endpoint Titer (1/n)
$10^6$
$10^5$
$10^4$
$10^3$
$10^2$
$10^1$
Prebleeds
Tri - 1Wk
Tri - 3Wk
MVA - 3Wk
B
NT50
$10^5$
$10^4$
$10^3$
$10^2$
$10^1$
$10^0$
Prebleeds
Tri - 1Wk
Tri - 3Wk
MVA - 3Wk

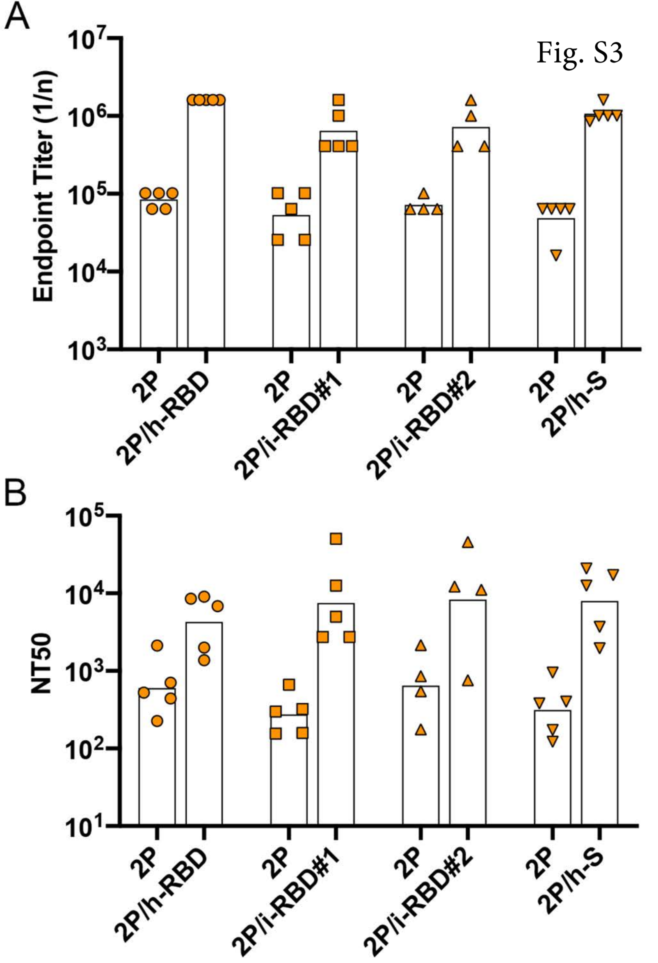
Fig. S3
A
B
Endpoint Titer (1/n)
$10^3$
$10^4$
$10^5$
$10^6$
$10^7$
NT50
$10^1$
$10^2$
$10^3$
$10^4$
$10^5$
2P
2P/h-RBD
2P
2P/i-RBD#1
2P
2P/i-RBD#2
2P
2P/h-S

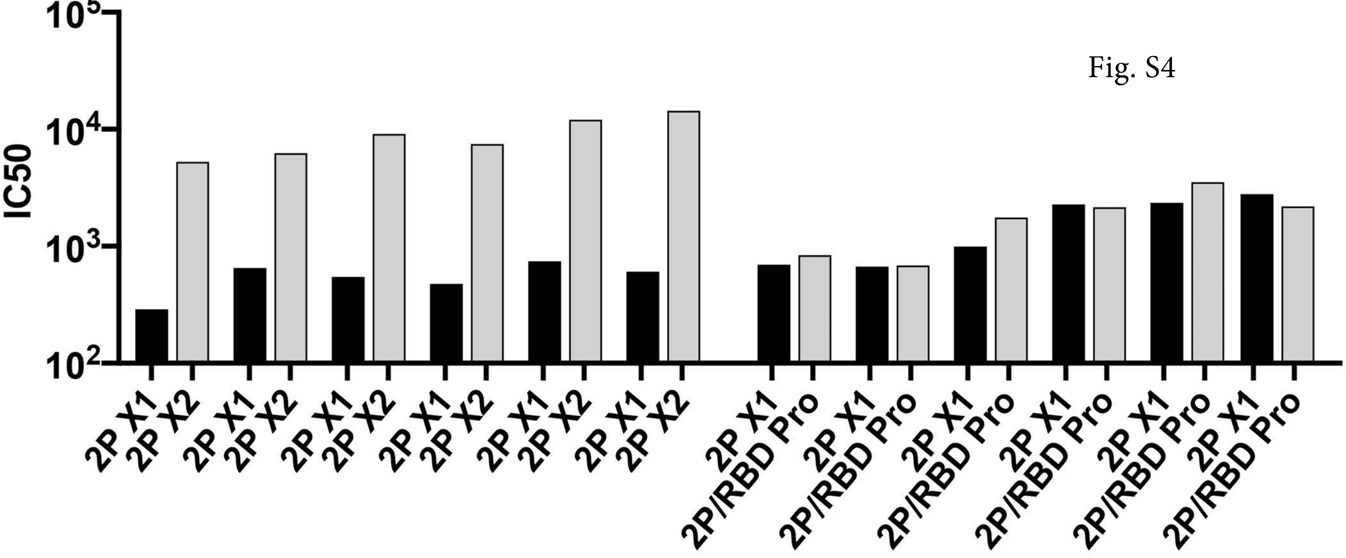
IC50
10^5
10^4
10^3
10^2
2P X1
2P X2
2P X1
2P X2
2P X1
2P X2
2P X1
2P X2
2P X1
2P X2
2P X1
2P X2
2P X1
2P/RBD Pro
2P X1
2P/RBD Pro
2P X1
2P/RBD Pro
2P X1
2P/RBD Pro
2P X1
2P/RBD Pro
2P X1
2P/RBD Pro

Fig. S4
